# Detection of *Plasmodium vivax* in a liver sample of a howler-monkey: one evidence more in favour of the identity between *Plasmodium simium* and *P. vivax*

**DOI:** 10.1101/2020.09.06.279869

**Authors:** J.C. Buery, A.M.R.C. Duarte, F.E.C. Alencar, C. Furieri, S.L. Mendes, A.C. Loss, C.R. Vicente, B. Fux, H.R. Rezende, P.V. Cravo, M.M. Medeiros, A.P. Arez, C. Cerutti

## Abstract

**Introduction:** The residual malaria of Atlantic Forest systems in Brazil occurs as an endemic disease with low frequency of cases. The chronological and spatial distance among the cases indicate an absence of fitness to the classical malaria cycle. This peculiar condition raised the suspicion of a reservoir, possibly the non-human primates. Simian and human malaria occur at the same places in that region, and there is already evidence of molecular identity between the simian parasites, *Plasmodium simium* and *Plasmodium brasilianum*, and the human parasites, *Plasmodium vivax* and *Plasmodium malariae*, respectively. Two different SNPs identified in the COX1 region of the *Plasmodium vivax/simium* of the Atlantic Forest reinforced its characterization as a zoonotic parasite. This finding supported the development of a PCR-RFLP protocol to identify such polymorphisms, and to monitor zoonotic malaria transmission.

**Methods:** In the present work, we tested the above-mentioned PCR-RFLP protocol in unprecedented mosquitoes and simian samples collected in Espírito Santo State, Brazil (ES).

**Results:** The parasite found in the simian sample was *P. vivax*, contrary to what the protocol should indicate. In the mosquito samples, the protocol disclosed both forms of the parasite.

**Conclusion:** This result suggests that the previously published pair of SNPs, and, consequently, the PCR-RFLP protocol, are not able to distinguish the dynamics of *Plasmodium* spp. circulation in the Atlantic Forest endemic area of ES.

## INTRODUCTION

Malaria is the most common parasitic disease worldwide, with 228 million cases and 405,000 deaths reported in 2018 (1). Six different species of *Plasmodium* are the etiologic agents for this mosquito-borne disease in humans (2). Global efforts aim to reduce the prevalence and mortality of malaria by 90% by 2030 in comparison to 2015 (212 million cases and 429,000 deaths globally) (3). However, several factors are hampering disease control, such as the presence of asymptomatic human carriers, drug and insecticide resistance, and, in particular cases, the existence of animal reservoirs that are often underestimated. The limited ability to identify asymptomatic carriers impairs malaria control, since these cases are not treated and thus remain as silent reservoirs for transmission. Furthermore, in specific endemic scenarios where non-human primates might participate in the life cycle of the parasite, the risk of ongoing transmission may also increase (4,5).

In the American continent, Brazil is the second country with the highest malaria frequency, with 194,271 cases in 2018, where *Plasmodium vivax, Plasmodium falciparum* and *Plasmodium malariae* are the most common etiologic agents. Most of the infections are caused by *P. vivax*, with the Amazon region comprising 99% of the reported cases nationwide. The remaining 1% transmission occurs in the extra-Amazonian Brazilian area. Although transmission in such areas was interrupted in the 1960s, there are still sporadic cases in regions of dense Atlantic Forest, with frequent case reports characterized by a special transmission cycle and clinical presentations. This unusual epidemiological setting has been termed *Bromeliad-malaria*, because the malaria-transmitting mosquitoes breed in the axils of the Bromeliaceae, being relatively frequent in the rural areas of the country’s coastal states. In such areas, *P. vivax* is commonly recognized as the etiologic agent of human infections, despite the occasional occurrence of *P. malariae*.

The Espírito Santo State (ES), located in the south eastern region of Brazil, is a model site to investigate the unusual transmission of *Bromeliad-malaria*, as it has large fragments of dense Atlantic Forest, and the highest number of cases nationwide. In what concerns the spatial and temporal distances between reported cases in these regions, the transmission cycle does not fit the traditional malaria cycle, as cases in nearby areas occur over a large time span between one and another, separated by several weeks, suggesting the occurrence of a zoonosis, with infected simians participating in the epidemiology. *Plasmodium simium* and *Plasmodium brasilianum* are the confirmed etiologic agents of simian infections, despite the studies based on genetic similarities suggesting that they are the same species as *P. vivax* and *P. malariae*, respectively (6-10).

Brasil *et al* (2017) (11) have previously studied a similar kind of transmission cycle in Rio de Janeiro State (RJ), by sequencing the complete mitochondrial genome (mtDNA) of *P. vivax/simium* recovered from humans and non-human primates. They detected a pair of SNPs in the COX1 gene, which was constant in all the samples tested, but different from sequences obtained previously from the Amazon region (11). While the sequences from RJ comprised the nucleotides C at position 3535 and G in the position 3869 those from the Amazon had, respectively, the nucleotides T and A at the corresponding positions. As a conclusion, the authors suggested that such SNPs are able to distinguish *P. simium*, classified as the simian parasite causing zoonotic malaria in Atlantic Forest, from *P. vivax*, the human parasite circulating in Amazon region. Buery *et al* (2017) (12) also sequenced the mitochondrial *P. vivax/simium* genome from diverse samples collected in an area of preserved Atlantic Forest in ES without migration history to the endemic states of the Amazon region and obtained different conclusions. Opposing the observations in RJ (11), the sequences obtained from *Anopheles* mosquitoes captured in the inner Atlantic Forest of ES revealed haplotypes consistent with the Amazon *P. vivax* SNPs pattern (12). Also, there was no common pattern of SNPs that allowed this classification of parasites in at least three samples from humans, meaning that these SNPs were not able to distinguish between the two forms in ES. Both studies importantly documented the presence of the same parasite species in both human and simian hosts. However, such findings are insufficient to confirm a zoonotic cycle.

A subsequent study by de Alvarenga *et al* (2018) (13) reported a novel PCR-RFLP protocol to detect polymorphisms in the mitochondrial COX1 gene, as previously proposed by Brasil *et al* (2017) (11). This protocol is directed at identifying *P. simium* in non-human primates and humans from the Atlantic Forest region, aiming to monitor zoonotic malaria transmission in Brazil. This was further used for epidemiological surveys in human and simian samples obtained in other Brazilian studies conducted in RJ (14, 15, 16). In a bid to test this simple and cost-effective tool in a different geographic area, our group used unprecedent ES samples, whose results we report herein.

## METHODS

In 2016, mosquitoes were collected in Valsugana Velha (VV), a district of Santa Teresa city (ES State), that is largely covered by well-preserved Atlantic Forest and where most of the *Bromeliad-malaria* takes place (19°57’58.4 “S, 40°34’45.2” W) (4). *Anopheles (Kerteszia) cruzii* female adults were pooled in groups of 10, which were collected in the same trap on the same date. Liver samples came from simians that perished due to yellow fever epizootics of 2016-2017 not far from the same region of Atlantic Forest fragments cited above. Sampling of simian and mosquito specimens was approved by the *Instituto Brasileiro do Meio Ambiente e dos Recursos Naturais Renováveis*/*Sistema de Autorização e Informação em Biodiversidade* (IBAMA/SISBIO) (licenses numbers 15.191 and 19.227-1, respectively). After DNA extraction, samples were screened for *P. vivax* trough a 18S rRNA based qPCR protocol (17). We then followed to characterize these samples according to the previously published protocol by de Alvarenga *et al* (2018) (13), followed by Sanger sequencing of the amplified amplicon, with minor adaptations.

Targeting the 18S rRNA gene, 29 samples were analysed, of which 16 were positive for *P. vivax*, corresponding to 14 pools of *A. K. cruzii* collected in tree canopies and two livers from monkey bodies. The proposed protocol to differentiate samples between *P. vivax* and *P. simium* (13) were successful for seven samples, six resulting from DNA pools of *A. K. cruzii* and one from a simian liver.

## RESULTS

Interestingly, *Plasmodium* DNA from the simian liver (sample F59) revealed to be *P. vivax*, in contrast to what could be expected in agreement with the previous reports by de Alvarenga *et al* (2018) (13). Regarding the mosquitoes, both *P. vivax* and *P. simium* were verified (Figure 1).

**Figure 1.**
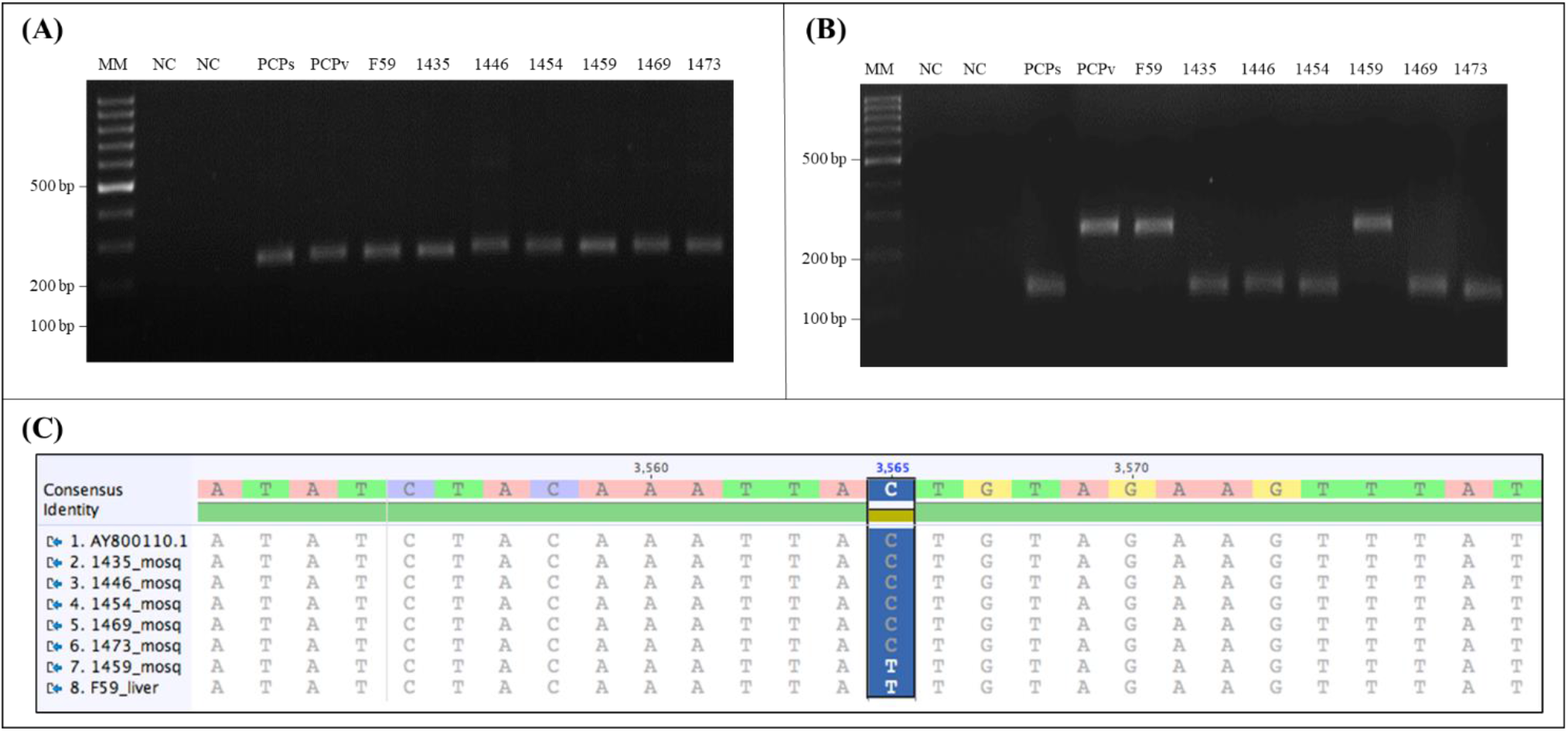
Proposed protocol of differential diagnosis of *P. simium* infection. **(A)** Nested/PCR followed by a **(B)** digestion with *Hpy*CH4III restriction enzyme of one simian liver (F59) and six mosquitoes (1435, 1446, 1454, 1459, 1469, 1473) samples. 3% agarose gel stained with GreenSafe. *Plasmodium vivax* DNA fragments are not digested (244bp), as *P. simium* DNA does, generating fragments of 118bp and 126bp. MM:1 kb Ladder. NC: Negative Control (no DNA). PCPs: positive control for *P. simium*, PCPv: Positive Control for *P. vivax*. **(C)** Alignment of partial mitochondrial COX1 gene sequence of *P. simium* isolated from free living simian liver (F59) from Atlantic forest; mosquito samples obtained in Atlantic Forest infected with *P. vivax* (1459) and *P. simium* (1435, 1446, 1454, 1469, 1473). GenBank sequence from *P. simium* (accession number at GenBank included in the name of sequence). The *Hpy*CH4III restriction site (ACNGT), including SNP T>C at position 3565, is highlighted.

## DISCUSSION

Lately, in studies performed in the Atlantic Forest region some authors classified all simian samples as *P. simium* under the de Alvarenga *et al* (2018) protocol indications (11, 13-16). Nevertheless, in ES, the parasite found in the simian host was classified as *P. vivax* by the same protocol, raising new issues about the classification of parasites in Atlantic Forest. This result suggests that the previously published SNPs and consequently the protocol to differentiate the parasites are not able to distinguish the dynamics of *Plasmodium* spp. circulation in the Atlantic Forest. When considering mosquito isolates and the interpretation of the above-mentioned protocol, the findings presented here suggest that both *P. vivax* and *P. simum* are likely to circulate on the canopy, with an equal probability of infecting simian hosts. At this point, the results show an updated scenario in the ES *Bromeliad-malaria* transmission. In 2017, Buery *et al* (12) reported a simian isolate from ES with the SNPs proposed by Brasil *et al* (2017) (11) as a *P. simium* based on the mtDNA sequence. Plus, mosquito isolates were analysed and none of them presented the nucleotides indicating the simian parasites. Recently, one sample of *Plasmodium* spp. DNA obtained from simian blood collected in southern Brazil was analysed by the same protocol and revealed the genetic pattern of *P. vivax* presented in the simian sample here reported (14). Unexpectedly, the authors designated the parasite as a non-*Plasmodium simium* (NPs, animal 4). Herein, the parasite disclosing this pattern was identical to *P. vivax* after qPCR, nested PCR, and Sanger sequencing. Taken together, these pieces of evidence reveal the genetic diversity in this heterogeneous and unusual cycle of transmission, and that only two SNPs do not seem to be sufficient to disclose the real epidemiological situation at Atlantic Forest region.

Improved understanding of the *Bromeliad-malaria* transmission cycle would strengthen the capability of Health authorities to promote adequate control strategies and would provide useful information to other Public Health authorities dealing with similar ecologic conditions around the world. The lack of a comprehensive understanding of the dynamics of the parasite’s transmission cycle in the Atlantic Forest is a consequence of a low number of both mammalian and vector samples which are analysed simultaneously but that are often collected in different periods. Accurate characterization of the distinctive genetic features of *Plasmodium* variants involved in *Bromeliad-malaria* requires the evaluation of all the putative elements of the cycle obtained from a cross-sectional survey or in a shorter timeframe. To verify a zoonotic cycle, one would have to analyse parasite genetic diversity and estimate the divergence time from the most recent common ancestor by molecular clock analysis. Such an approach was performed to analyse the transmission of *P. knowlesi*, uncovering a higher number of genotypes per infection in simians than in humans (18), which allowed the first unequivocal confirmation of zoonotic transmission of malaria in Southeast Asia (18). Similarly, natural transmission of *Plasmodium cynomolgi* between simians and humans became known in 2011 in Malaysia (19). Divergence time from the most recent common ancestor based on the analysis of mtDNA revealed the derivation of the species from an ancestral parasite population that existed before human settlement in Southeast Asia. Both findings were able to support the hypothesis of an actual zoonosis, pointing to a recent transfer of the parasites to the humans. In the case of *P. vivax/simium*, however, the evidence points in the opposite direction. The haplotype diversity is lower among the simians, and the phylogenetic analyses indicate a recent transfer of the species from humans to simians (12, 20).

## CONCLUSION

The results of this work suggest that the published SNPs and the protocol of differential diagnosis of *P. simium* infection are not able to elucidate the dynamics of *Plasmodium* spp. circulation in the Atlantic Forest of ES. The evidence until now is not sufficient to determine how and if the transfer of the parasites occurs in the Atlantic Forest, precluding any definite conclusion regarding a zoonotic cycle. Further studies with larger numbers of samples from humans, simians, and vectors, collected over a short timeframe will allow deeper understanding of the transmission cycle of this singular endemic disease.

## Author contributions

***Buery, J.C***.: Conceptualization, Validation, Investigation, Writing - original draft, Project administration, Funding acquisition. ***Duarte, A.M.R.C***.: Resources, Investigation, Writing - Review & Editing. ***Alencar, F.E.C. and Vicente, C.R***.: Writing - Review & Editing, Visualization. ***Furieri, C*.; *Mendes, S.L*.; *Rezende, H.R. and Fux, B***.: Resources, Data Curation. ***Loss, A.C***.: Formal analysis, Writing - Review & Editing. ***Cravo, P.V. and Medeiros, M.M***.: Validation, Writing - Review & Editing. ***Arez, A.P***.: Validation, Resources, Writing - Review & Editing, Supervision. ***Cerutti Jr, C***.: Conceptualization, Resources, Writing - original draft, Project administration, Supervision.

